# Facial expressions of emotional stress in horses

**DOI:** 10.1101/2020.10.19.345231

**Authors:** Johan Lundblad, Maheen Rashid, Marie Rhodin, Pia Haubro Andersen

## Abstract

Horses have the ability to generate a remarkable repertoire of facial expressions, some which have been linked to certain emotional states, for example pain. Studies suggest that facial expressions may be a more ‘honest’ expression of emotional state in horses than behavioral or physiological parameters. This study sought to describe the facial expressions during stress of healthy horses free of pain, using a standardized method of recording facial expressions in video. Stress was induced in 28 horses by subjecting them to road transport and 10 of these horses were also subjected to social isolation. The horses served as their own control. A body-mounted, remote controlled heart rate monitor provided continuous heart rate measurements during the interventions. The horses’ facial expressions were video-recorded during the interventions. Frequency and duration of each facial expression were then determined, according to the Equine Facial Action Coding System. Heart rate increased during the stressful interventions (p=0.01), confirming that the interventions were stressful. Using both the human investigation- and the co-occurrence methods, the following facial traits could be observed during stress: *eye white increase* (p<0.001), *nostril dilator* (p<0.001), *upper eyelid raiser* (p<0.001), *inner brow raiser* (p=0.042), *tongue show* (p<0.001) along with an increase in ‘ear flicker’ (p<0.001) and blink frequency (p<0.001). The facial actions were successfully used to train a machine-learning classifier to discriminate between stressed and calm horses, with an accuracy of 74.2 %. Most of the facial features identified correspond well with previous research on the subject, for example flared nostrils, repetitive mouth behaviors, increased eye white, tongue show and ear movements. Some features selected as indicative of emotional pain-free stress are used in face-based pain assessment tools, such as dilated nostrils, eye white increase or inner brow raiser. The relation between facial expressions of stress and pain should therefore further be studied.

## Introduction

Lack of a ‘gold standard’ for evaluating emotional states in non-verbal mammals has been a driving force for exploration of bodily behavior or physiological markers to convey information about internal states [1]. Facial activity can generate a wide array of different observable expressions [2] and has been suggested as a tool for assessment of welfare in mammals [1]. Many facial expressions are conserved across mammal species, including humans [3]. In humans, who can self-report, it is known that the affective component of pain is expressed by prototypical facial expressions [4]. The human emotional states happiness, fear, anger and disgust are also associated with typical facial expressions [5]. Horses have the ability to generate a remarkable repertoire of facial expressions, which can be described by 17 action units [6]. This is smaller than the human repertoire [7], but larger than that of e.g., chimpanzees or dogs [8,9]. Recently, it has been shown that horses can display facial changes which are specific to pain [10–12]. The facial actions involved include features such as eyebrow raising or tightening of eyelids, ears back, tension of the lower face muscles, and widened nostrils. Current pain assessment tools use these traits [10–12], but with some differences in the descriptions of facial configurations.

One limitation to using facial cues for pain assessment in horses is that the specificity in relation to other common affective states, e.g., emotional stress, is not known [13]. Stress induces typical facial expressions in humans [14], but only a few studies of facial expressions during stress have been performed in horses, most focusing on features around the eye [15] or blinking frequency to determine stress. However, there is some controversy on whether the frequency of blinks increases [16] or diminishes [17] during stress. Pain is an internal stressor and the experience of pain may therefore elicit a stress response. In contrast, a stress response does not infer pain. From human research, it is known that emotional state in individuals affects their perception of pain [5]. These complex interactions between stress and pain are currently unresolved in horses and other animals [6].

Stress is an adaptive physiological and emotional response that enables coping with challenges from the environment, such as tissue damage, or perceived threats of injury. In horses, which are prey animals, a multitude of emotional challenges may be present during both ordinary and extraordinary situations. These situations may include competitions, transportation by road, separation from the herd, social isolation during transportation, introduction to a new environment, and exposure to confinement during diagnostic procedures and treatment at an equine hospital. Most of these handling procedures have been shown to induce emotional stress [18–20], in addition to the stress inevitably induced by any pain the horse may feel at the moment.

The stress response may clinically appear as elevated heart and respiratory rate, increased blood pressure, and temperature [21]. It may even induce some degree of analgesia [22] or hyperalgesia [23], at least experimentally. These responses occur together with a number of behavioral changes, such as alertness or aversiveness (short-term) and stereotypies or apathy (long-term) in horses [21]. Some stress-related physiological changes are not specific to emotional stress, however. Cortisol release has a diurnal variation [24] and may be affected by pathologies [25]. Heart rate and blood pressure may be elevated in response to both positive and negative experiences, such as exercise [26], or during experience of an internal stressor, such as pain [25]. This renders physiological markers suitable for measurements of emotional stress in controlled settings, but not in the field, where discrimination between stress and pain is important in clinical decision making. It is therefore of interest to investigate whether the rich facial repertoire of horses contains distinct facial expressions of emotional stress.

The Equine Facial Action Coding System (EquiFACS) [6] is a tool for recording facial expressions by observing onset and offset of anatomically based action units (AUs) and action descriptors (ADs) over time. The method does not infer anything about the meaning of the observed facial movements, leaving less space for subjective judgment. The resulting dataset contains data on the occurrence of different action units, time of onset, offset, and duration, and their temporal overlap with other active action units. EquiFACS was used to determine facial expressions of pain only recently [27]. To determine actions that are typical for pain in humans, a method based on increased frequencies is commonly used [28]. However, statistical methods for analyzing FACS data on horses are not yet well-developed, and data-driven methods for classification of emotional state in horses based on the frequency and duration of action units are generally lacking.

The aim of this study was therefore to describe, using EquiFACS, the facial expressions during controlled stressful events in healthy horses free of pain. Based on earlier descriptions, we expected to identify facial action patterns that were distinct to stress, with the most prominent being changes in repetitive mouth behaviors, flared nostrils, flattened ears [29], and the action descriptors yawning and tongue show [30]. We also expected an increased number of action units in response to visual or auditory inputs, displayed as increased frequencies of ear movement and eye blinks. To our knowledge, facial expressions of emotional stress have not been described previously using EquiFACS.

Short-term emotional stress was induced by transporting healthy horses in a trailer or keeping them in short-term social isolation. The physiological parameter heart rate was used as a marker of induced stress. We hypothesized that the frequency methods applied in human research can also identify important action units and action descriptors in horses, but also that methods using temporal distribution are accurate in classification of stress in horses.

## Materials and methods

### Ethical statement

This study was approved by the Ethics Committee for Animal Experiments, “Blinded for review” (Approval no. 5.8.18-10767/2019). Owner consent for use of privately owned horses was obtained before experimentation.

### Study design

For this observational study, two standard horse management practices were used to induce emotional stress: short-term transportation and short-term isolation. Video footage was recorded during the stressful events, and when the horse was calm before or after the intervention. A body-mounted, remote-controlled heart rate monitor provided continuous heart rate measurements in all three situations.

### Horses

A total of 28 university-owned and privately-owned horses were used in the study. Nine Standardbred trotters (seven mares and two geldings), and one warmblood mare from the university herd (UNI) were included. They were considered healthy at routine examinations during the previous four months, were of median age 12 years (range 8-19 years), and had roughly the same body weight. They were kept at an authorized research facility at “Blinded for Review”. These horses were fed hay four times a day, and oats once a day according to a nutritional plan that supported normal condition. All horses were allowed out on pasture for 6 hours a day and otherwise kept in individual 3 m x 4 m boxes. During the experiment, horses were moved to other boxes in the same facility and acclimatized for at least 16 hours. Horses were moved together in pairs, stabled besides each other, and kept in their regular stable herd (together for at least the previous 6 months). Each pair of horses had the same feeding and housing routine and had the same caretakers in all stables.

Eighteen privately owned horses (PRI) were included. They comprised 10 geldings, seven mares and one stallion, of the breeds Thoroughbreds (n=5), mixed-breed ponies (n=4), Standardbred trotters (n=3), and Swedish warmblood/riding breeds (n=6), with body weight ranging between approximately 400 and 600 kg. The median age of horses in this group was 10 years (range 3-24 years). They were considered healthy by their caretakers, had not been subjected to veterinary treatment for the previous two months, and had not been treated with analgesics during that period. The horses were managed at home, by the horse owner, in the routines to which they were accustomed. All were kept in stables except for the thoroughbreds, which were kept in a free-range system. Three of the PRI horses were kept at the university but were treated as though they were privately owned.

All horses from PRI (N=18) and UNI (N=10) were subject to transportation (N=28). The PRI horses were studied in their own stable and were transported in their own trailer. The UNI horses were transported in a standard horse trailer for 20 minutes. All horses from UNI were subjected to social isolation on a subsequent occasion (N=10). Social isolation was performed by leaving the horse alone in the stable without its herd mates for at least 15 minutes.

### Video-recording

Video-recordings of the horses were made during the two types of stress and during baseline without the presence of an observer. In the PRI group and during transportation in the UNI group, video-recordings were made in the box and inside the horse trailer, using GoPro Hero 3+ Silver Edition and GoPro Hero 7 Black cameras (Gopro Inc., San Mateo, California, USA). Resolution was set to 1080p at 30 fps and videos were exported to mp4-format. The cameras were mounted depending on the layout of the box, so that the entire horse and its box could be seen in the footage. If the stable had no regular box, the horses were filmed in their grooming spot. In the trailer, the halter of the horse was tied to a front bar in a standard manner, and the camera was mounted in line with the horses’ head height and angled approximately 45-60 degrees from the horses’ medial plane. The cameras recorded for 10 to 20 minutes during transportation, and for at least 30 minutes during baseline.

During the social isolation intervention and the baseline (UNI), the horses were filmed in their own boxes. The video-recordings during social isolation were made using two wall-mounted standard surveillance cameras with night vision (WDR EXIR Turret Network Camera, HIKVISION, Hangzhou, China). Extra light was provided with nine standard fluorescent lights mounted in the ceiling, programmed to provide light during daytime hours. The cameras were mounted in each corner in the front of the box so only the horse and its box could be seen in the footage, in order to ensure blinding. Resolution was set to maximum and images were exported to mp4-format. The cameras recorded all baseline sessions for a minimum of 30 minutes and social isolation sessions for a minimum of 15 minutes.

### Heart rate monitoring

A remotely controlled human sport ECG transmitter (Polar Wearlink, Polar Electro OY, Kempele, Finland), modified for equine use, was used together with its corresponding control unit to obtain continuous heart rate measurements without the interference of an observer. The Wearlink device was fastened using a girth, which was soaked in water before attachment. The horses were allowed to adjust to the ECG transmitter for 10 minutes before measurements began [31]. The heart rate monitor was synchronized with the cameras, using a gesture in the video when the transmitter was started or using the time-stamped files produced by the cameras and heart rate transmitter. Files containing R-R intervals were exported and filtered through Polar ProTrainer Equine Edition (Polar Electro OY, Kempele, Finland). The files were processed in Kubios HRV Premium (Kubios, Finland) in order to extract heart rate during the selected time intervals. Heart rate measurements were extracted as a mean during five minutes, with onset two minutes and 15 seconds before annotation started. A two-way paired t-test for means was used to calculate significance in the PRI group. In the UNI group, ANOVA was used to test for significance between all three interventions and a two-way paired t-test for means was used to test for the specific rise in heart rate between the baseline and the respective intervention.

### Video processing and annotation

The identity of the video-recordings of the transportation group could only be blinded for horse, and not for intervention, since the location in the trailer and its movements could not be hidden. Selection of clips was made by manual inspection and 30-second clips of suitable footage were cut from the videos. If the face was visible and scorable for more than 30 seconds, a random number generator was used for video selection.

The identity of the video-recordings of the social isolation group was blinded in relation to horse and intervention before annotation. Selection of videos for the social isolation group was performed using an automated face detection system [32], where sequences were selected if the head position of the horse was suited for annotation. Thirty-second sequences of video with a side- or front-view confidence of at least 60% were selected. If several selections were available, a random number generator was used to select one clip. The selected clips were manually inspected to ensure that the software had successfully identified a face. If not, a new clip was randomly selected.

All films were annotated in a blinded manner by two EquiFACS-certified veterinarians with a minimum of 70% correct annotations compared with expert raters. All transportation and baseline films were also annotated by JL who is also certified in EquiFACS. Annotation was performed using a template consisting of all codes in EquiFACS, including supplemental codes and the visibility code VC74 (code for unscorable), but without head movements (AD51-AD55). Annotation was performed with the open-source program ELAN [33]. The annotators coded the onset and offset of the facial action units, allowing calculation of frequency and duration, i.e., how frequently an action unit or action descriptor occurred and how long it remained active. The annotators set the onset of the action unit to when the muscle started contraction and the offset to when it was fully back to neutral again. Inter-rater agreement between the coders was calculated using the Wexler ratio as described by Ekman et al. [7], using a full 30-second clip as the sample. Inter-rater agreement was calculated to be on average 0.75 (coder 1-2: 0.76; coder 2-3: 0.76; coder 1-3: 0.71), indicating good agreement between raters.

### Selection of EquiFACS codes in stressed horses

Since inter-rater agreement was good, one set of annotations was randomly selected and used for each video. For each selected action unit or action descriptor, frequency and duration were observed. Frequency of ear flicker movements was also investigated. In order to do this, a movement index was created, by describing *ears forward* (EAD101) and *ear rotator* (EAD104) occurring together within a one-second interval. It is important to note that this is not an action descriptor but a definition of an occurrence, where the selected action descriptors occur in succession to constitute the “ear flicker”.

EquiFACS codes were analyzed using the method described by Kunz et al. [28], here called the Human FACS Investigation (HFI) method. Action units that accounted for more than 5% of total action unit occurrences in stress videos were selected. From this subset, action units detected at higher frequency in stress videos than in no-stress videos were selected as the final set of stress action units. While the HFI action unit selection method ensures that selected codes are frequent and distinct, they may have only a slightly stronger correlation with the emotional state and can exclude less frequent, but highly discriminative, action units. Therefore, the relative temporal distribution of action units was also considered. In order to do this, the method of Rashid et al. [27], here referred to as the Co-occurrence method, was used to calculate the co-occurrence of action units. This method selected EquiFACS codes that occurred together with other EquiFACS codes more frequently in stress than in no-stress states. Since onset and offset of EquiFACS codes were recorded in ELAN, codes which appeared simultaneously or in close relation to each other could be further studied. EquiFACS codes that occurred within a predetermined period (observation window size, OWS) were recorded as co-occurring. Action units that exhibited the largest difference in co-occurrence patterns between stress and no-stress states were selected. The method uses directed graphs to record and calculate differences in co-occurrence patterns. Furthermore, a paired t-test for mean values was used to test significance, with p<0.05 considered significant.

For both the HFI and Co-occurrence methods, occurrences of *ears forward* (EAD101) and *ear rotator* (EAD104) that were included in the “ear flicker” category were not double counted for EAD101 and EAD104 separately. As a result, occurrence counts of EAD101 and EAD104 did not occur within a one-second interval of one another.

### Classification of stress/no stress

The EquiFACS codes selected by the HFI and Co-Occurrence methods were used to train a machine learning classifier, Linear Support Vector Machine (LVSM), for stress versus no-stress classification. Twenty-five baseline videos and 35 stress videos (10 from social isolation, 25 from transportation) were used. The frequency and duration features in the clips were used to represent each video sequence, in order to train the LSVM for stress versus no-stress classification. Using five-fold cross-validation, the optimum regularization parameter *C* and balanced class weights were selected. The Python Scikit-Learn library [34] and the Leave-One-Out (LOO) protocol were used to train and test the models, meaning that the features of all videos except one were used to train an LSVM that then used the same features on the remaining video to determine whether it showed a stress state. The LSVM predictions were collated across the entire dataset, and precision, recall, and accuracy were computed. The performance of the LSVM models indicated how well the selected EquiFACS codes captured the facial expressions of stress and acted as a type of construct validity to classify stress.

## Results

### Heart rate during interventions

The heart rate during interventions is shown in Fig 1. Heart rate increased from a pooled mean of 41 bpm (SD 10.6) during baseline to 70 bpm (SD 24.3) during transportation and to 55 bpm (SD 21.9) during social isolation. The increase in both groups was significant (p<0.01). In general, the spread of samples in the transportation group indicated that these measurements were somewhat more disrupted, due to more movement of the horse, but in general the heart rate samples were of good quality. Heart rate after the interventions decreased fully to the baseline level, indicating that it was the intervention that caused the rise in heart rate. The data also indicated that the transportation intervention was more stressful to the horses than the social isolation intervention. Based on the similarities in these results, the PRI and UNI groups were regarded as one stress group in the following analysis of facial expressions.

**Fig 1.**
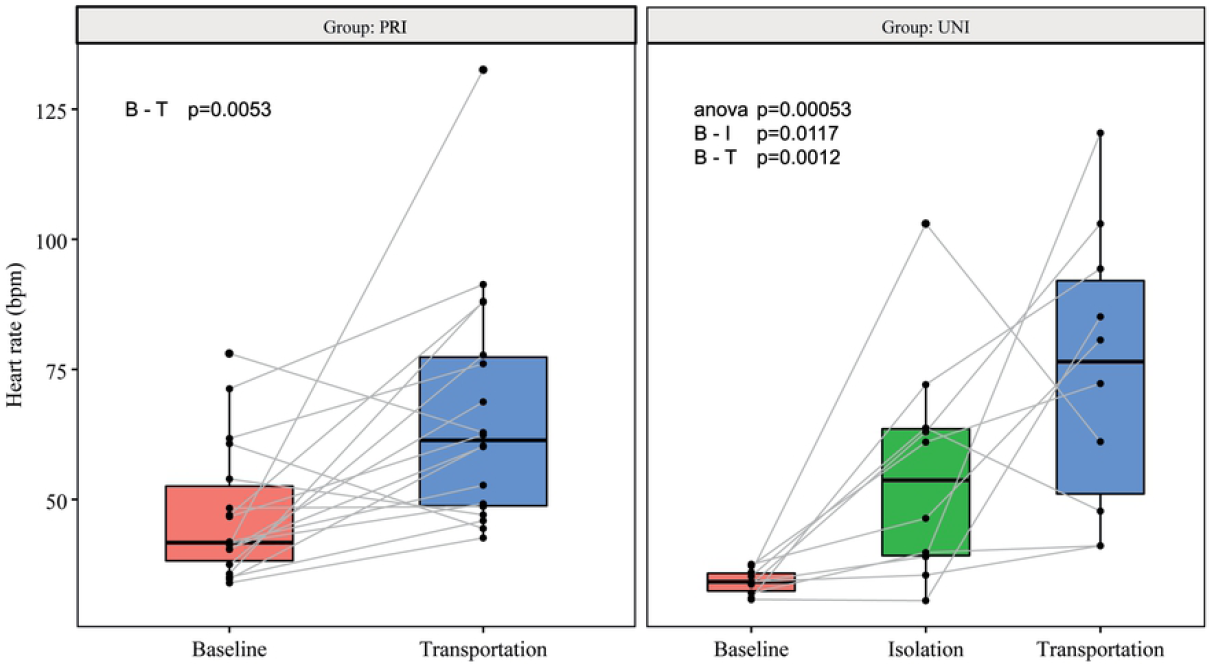
Heart rate during interventions. Boxplots showing the heart rate of (left) privately owned horses (PRI) and (right) university horses (UNI) during baseline (B), isolation (I), and transportation (T).

### Selected EquiFACS codes (HFI method)

Table 1 shows action units selected using the HFI method. All action units that comprised at least 5% of stress action unit occurrences were more frequent in transportation videos than baseline videos. Blink action units (AU145 and AU47) and *inner brow raiser* (AU101) had the most similar rate of occurrence between stress and no-stress states, while *eye white increase* (AD1), *nostril dilator* (AD38) and *upper lid raising* (AU5) exhibited the largest difference in frequency between transportation and baseline recordings. The selected action units for social isolation stress mostly showed similar codes. Unlike for the transportation group, *half blink* (AU47) was not selected for the social isolation group due to occurring more frequently in no-stress videos, and *upper lid raiser* (AU5) is not selected due to low frequency in social isolation videos. On the other hand, *ear rotator* (EAD104) was selected during social isolation. “Ear flicker” was more frequent and more pronounced in transportation than in social isolation.

**Table 1.** Facial expressions during stress (HFI method)

|  | *Eye white increase* (AD1) | *Nostril dilator* (AD38) | *Inner brow raiser* (AU101) | *Blink* (AU145) | *Half blink* (AU47) | *Upper lid raiser* (AU5) | “Ear flicker” | *Ear rotator* (EAD104) |
| --- | --- | --- | --- | --- | --- | --- | --- | --- |
| **Transportation** |  |  |  |  |  |  |  |  |
| **Percentage of AUs during stress / no stress** | 8.2% / 4.8% | 13.1% / 8.4% | 5.3% / 8.1% | 12.7% / 19.6% | 7.7% / 11.2% | 8.0% / 5.7% | 18.9 / 17.7% | Not selected |
| **Difference in frequency** | 113.7% | 106.2% | 31.4% | 30.1% | 35.8% | 98.6% | 76.2% | Not selected |
| **Isolation** |  |  |  |  |  |  |  |  |
| **Percentage of AUs during stress / no stress** | 7.8% / 3.9% | 15.0% / 7.0% | 15.0% / 12.1% | 18.1% / 19.8% | Not selected | Not selected | 16.6% / 20.2% | 5.3% / 6.2% |
| **Difference in frequency** | 85.7% | 90.9% | 43.0% | 12.8% | Not selected | Not selected | 1.9% | 6.1% |
| **Combined** |  |  |  |  |  |  |  |  |
| **Percentage of AUs during stress / no stress** | 8.2% / 4.8% | 13.4% / 8.4% | 7.2% / 8.1% | 13.8% / 19.6% | 8.0% / 11.2% | 7.0% / 5.7% | 18.4% / 17.7% | Not selected |

| Difference in frequency | 106.8% | 101.8% | 52.1% | 29.3% | 30.5% | 80.7% | 66.4% | Not selected |
| --- | --- | --- | --- | --- | --- | --- | --- | --- |

Action units (AUs) and action descriptors (ADs) selected using the Human FACS
Investigation (HFI) method to represent stress in horses in the transportation and social
isolation groups and together as a combined group.

The selected action units when both groups were combined were used as a larger ‘stress’ group. All action units that comprised at least 5% of stress action unit occurrences were also more frequent in stress videos than baseline videos. The chosen action units for social isolation were identical to those selected for transportation stress, but the percentage difference between no-stress and stress frequency counts was noticeably larger for *inner brow raiser* (AU101).

### Frequency and duration of AUs

Average frequency and maximum duration for selected action units across baseline, social isolation, and transportation videos are shown in Fig 2. Action unit frequency increased across all selected action units, particularly between baseline and transportation stress. However, not all rises were significant.

**Fig 2.**
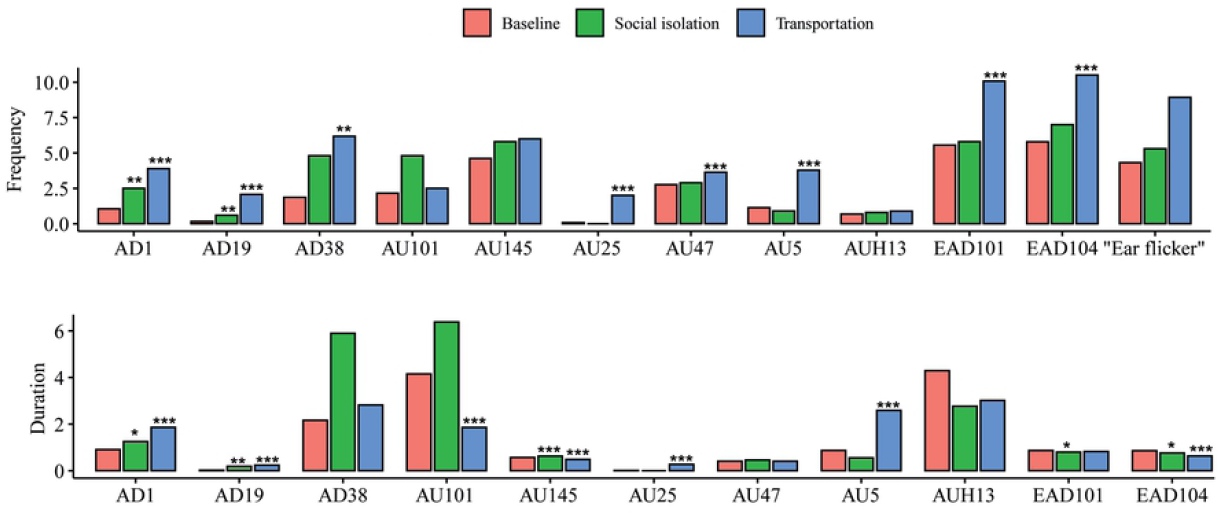
Frequency and duration of EquiFACS codes. Changes in action unit (AU) and action descriptor (AD) frequency patterns between stress and no-stress states. Stress affected the duration (s) of activity for an AU. Asterisks indicate statistical significance (*p<0.05, **p<0.01; ***p<0.001).

Stress, particularly transportation stress, was correlated with an increase in duration of *upper lid raiser* (AU5), *eye white increase* (AU101), *inner brow raiser* (AU 101), and *nostril dilator* (AD38). All action units selected by the HFI method had p<0.01 for at least one representation and group. Additionally, *tongue show* (AD19) and *lips part* (AU25), related to mouth behavior, showed p<0.01 across all groups and representations when tested separately. However, each of these action units occurred rarely. *Inner brow raiser* (AU101), despite its high frequency, was only statistically significant when using maximum duration and decreased during transport stress. With just 10 horses in the group, action unit frequency was rarely significant for isolation stress.

### Selected EquiFACS codes (Co-occurrence method)

Action units and action descriptors selected using the Co-occurrence method are presented in Table 2. Of the selected codes, *nostril dilator* (AD38), *tongue show* (AD19), *mouth open* (AU25), *upper lid raiser* (AU5), *eye white show* (AD1), and “ear flicker” showed significance in all OWS. *Inner brow raiser* (AU101) was selected by the HFI method and significant (up to a 5-second OWS) using this method.

**Table 2.**
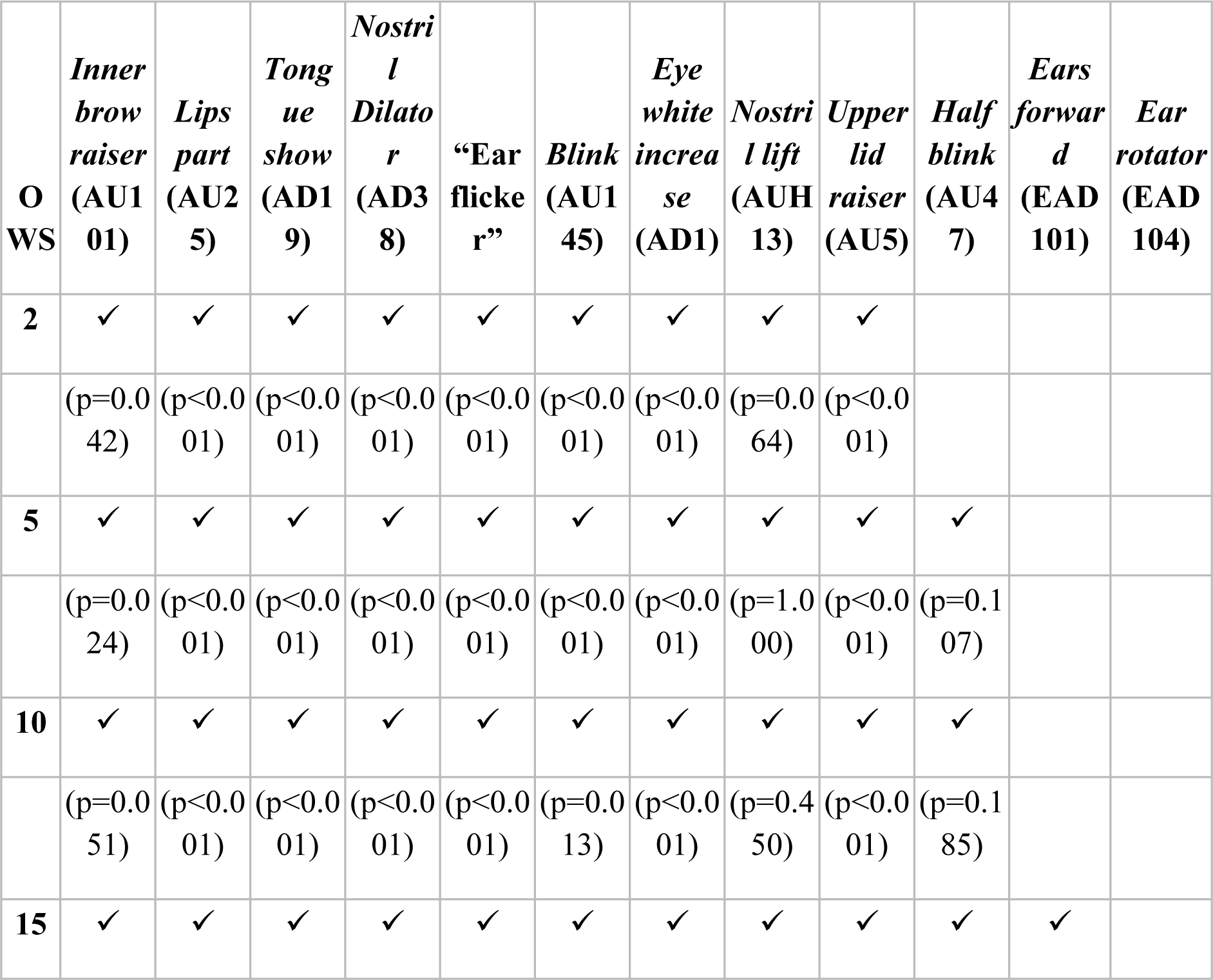
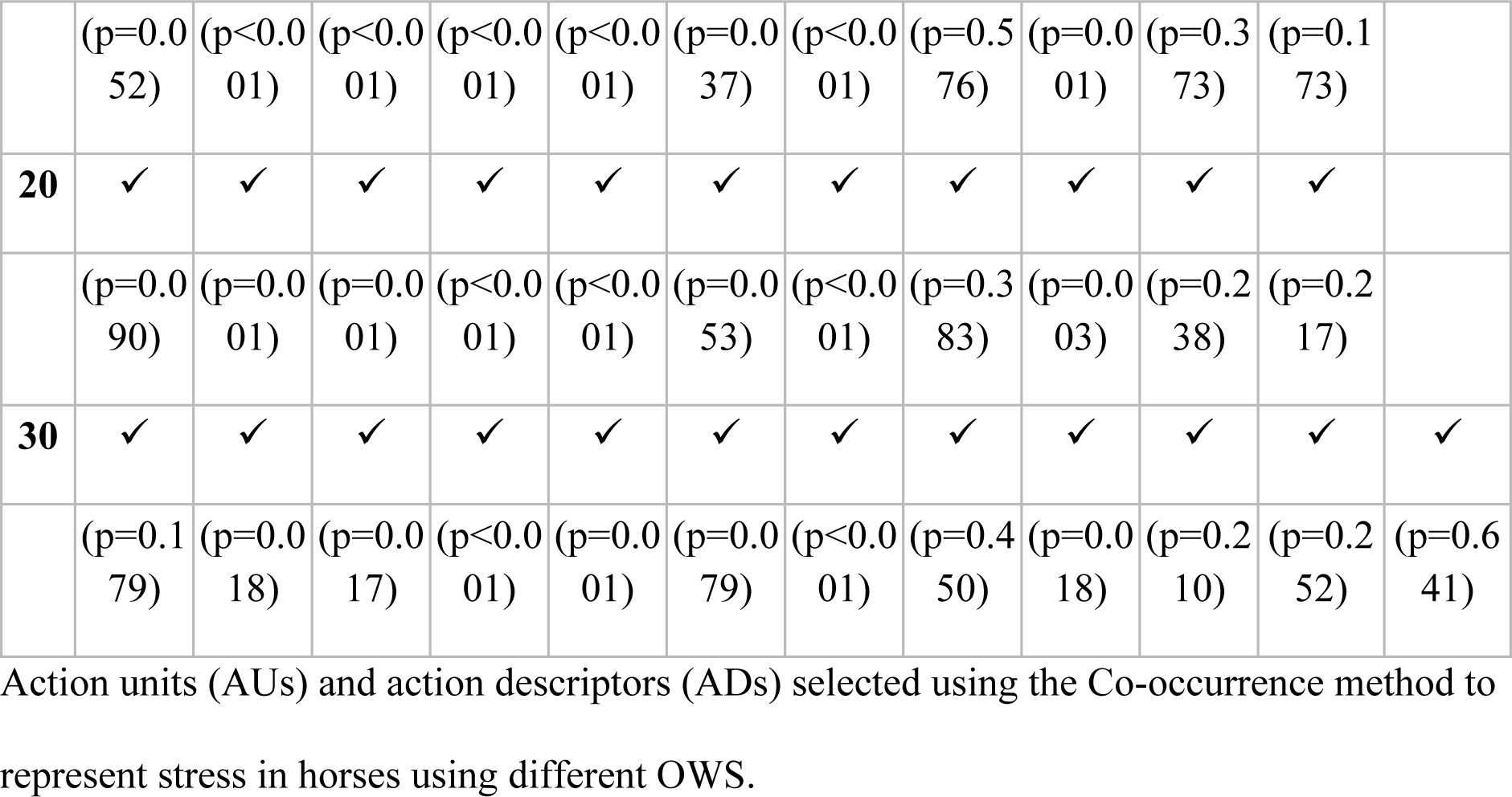
Facial expressions during stress (Co-oc method).

| **O**<br>**WS** | ***Inner***<br>***brow***<br>***raiser***<br>**(AU1**<br>**01)** | ***Lips***<br>***part***<br>**(AU2**<br>**5)** | ***Tong***<br>***ue***<br>***show***<br>**(AD1**<br>**9)** | ***Nostril***<br>***Dilato***<br>***r***<br>**(AD3**<br>**8)** | **“Ear**<br>**flicke**<br>**r”** | ***Blink***<br>**(AU1**<br>**45)** | ***Eye***<br>***white***<br>***increa***<br>***se***<br>**(AD1)** | ***Nostril***<br>***lift***<br>**(AUH**<br>**13)** | ***Upper***<br>***lid***<br>***raiser***<br>**(AU5)** | ***Half***<br>***blink***<br>**(AU4**<br>**7)** | ***Ears***<br>***forward***<br>**(EAD**<br>**101)** | ***Ear***<br>***rotator***<br>**(EAD**<br>**104)** |
| --- | --- | --- | --- | --- | --- | --- | --- | --- | --- | --- | --- | --- |
| **2** | ✓ | ✓ | ✓ | ✓ | ✓ | ✓ | ✓ | ✓ | ✓ |  |  |  |
|  | (p=0.0<br>42) | (p<0.0<br>01) | (p<0.0<br>01) | (p<0.0<br>01) | (p<0.0<br>01) | (p<0.0<br>01) | (p<0.0<br>01) | (p=0.0<br>64) | (p<0.0<br>01) |  |  |  |
| **5** | ✓ | ✓ | ✓ | ✓ | ✓ | ✓ | ✓ | ✓ | ✓ | ✓ |  |  |
|  | (p=0.0<br>24) | (p<0.0<br>01) | (p<0.0<br>01) | (p<0.0<br>01) | (p<0.0<br>01) | (p<0.0<br>01) | (p<0.0<br>01) | (p=1.0<br>00) | (p<0.0<br>01) | (p=0.1<br>07) |  |  |
| **10** | ✓ | ✓ | ✓ | ✓ | ✓ | ✓ | ✓ | ✓ | ✓ | ✓ |  |  |
|  | (p=0.0<br>51) | (p<0.0<br>01) | (p<0.0<br>01) | (p<0.0<br>01) | (p<0.0<br>01) | (p=0.0<br>13) | (p<0.0<br>01) | (p=0.4<br>50) | (p<0.0<br>01) | (p=0.1<br>85) |  |  |
| **15** | ✓ | ✓ | ✓ | ✓ | ✓ | ✓ | ✓ | ✓ | ✓ | ✓ | ✓ |  |
|  | (p=0.0<br>52) | (p<0.0<br>01) | (p<0.0<br>01) | (p<0.0<br>01) | (p<0.0<br>01) | (p=0.0<br>37) | (p<0.0<br>01) | (p=0.5<br>76) | (p=0.0<br>01) | (p=0.3<br>73) | (p=0.1<br>73) |  |
| **20** | ✓ | ✓ | ✓ | ✓ | ✓ | ✓ | ✓ | ✓ | ✓ | ✓ | ✓ |  |
|  | (p=0.0<br>90) | (p=0.0<br>01) | (p=0.0<br>01) | (p<0.0<br>01) | (p<0.0<br>01) | (p=0.0<br>53) | (p<0.0<br>01) | (p=0.3<br>83) | (p=0.0<br>03) | (p=0.2<br>38) | (p=0.2<br>17) |  |
| **30** | ✓ | ✓ | ✓ | ✓ | ✓ | ✓ | ✓ | ✓ | ✓ | ✓ | ✓ | ✓ |
|  | (p=0.1<br>79) | (p=0.0<br>18) | (p=0.0<br>17) | (p<0.0<br>01) | (p=0.0<br>01) | (p=0.0<br>79) | (p<0.0<br>01) | (p=0.4<br>50) | (p=0.0<br>18) | (p=0.2<br>10) | (p=0.2<br>52) | (p=0.6<br>41) |

Action units (AUs) and action descriptors (ADs) selected using the Co-occurrence method to represent stress in horses using different OWS.

### Leave-One-Out classification

The selected action units in Tables 1 and 2 were used to train a simple LVSM for stress or no-stress classification, in order to check the validity of the selected action units. Results of the LOO classification are shown in Tables 3 and 4. The best classification was obtained using both frequency and maximum duration, reaching an impressive 89% recall rate for the action units selected by the HFI method and a 78% precision rate for the action units selected by the Co-occurrence method.

**Table 3.** Results of Leave-One-Out classification for the action units selected by the Human FACS Investigation (HFI) method

|  | Frequency | Max duration | Both |
| --- | --- | --- | --- |
| **Precision** | 73.91% | 71.43% | 66.67% |
| **Recall** | 89.47% | 92.11% | 89.47% |
| **Accuracy** | 75.76% | 74.24% | 68.18% |

**Table 4.** Results of Leave-One-Out classification for the action units selected by the Co-occurrence method

|  | Frequency | Max duration | Both |
| --- | --- | --- | --- |
| **Precision** | 72.09% | 58.82% | 78.38% |
| **Recall** | 81.58% | 52.63% | 76.32% |
| **Accuracy** | 71.21% | 51.52% | 74.24% |

## Discussion

The basis for the emotional component of stress was disrupting the horses’ regular routines, by either keeping them in while their herd mate was brought outside or by loading them onto a trailer for transportation. However, it is possible that a number of external inputs inevitably associated with transportation, e.g., exposure to new environment, wind, confined space, social isolation, or movement restriction, induced additional stress. Social isolation, on the other hand, was associated with few external inputs, because the horses stayed in their familiar environment during the intervention. In the social isolation group, a significant rise in heart rate was observed, although to a lower level than during transportation. It was concluded that both interventions induced emotional stress in horses, according to the rise in heart rate [19,21,30,35,36] recorded under well-controlled circumstances.

Analysis of the EquiFACS codes showed increased frequency of several specific action units and action descriptors during both interventions (Fig 3). According to the HFI method, the action units of a stressed horse included *upper lid raiser* (AU5) and *inner brow raiser* (AU101), as well as *blink* (AU145) and “ear flicker”. The frequency of the action descriptors *nostril dilator* (AD38) and *eye white increase* (AD1), not describing certain muscle-induced codes but rather the effects of other muscle movement, was also significantly increased. According to the Co-occurrence method, *tongue show* (AD19) and *mouth open* (AU25) were also important. When comparing the HFI method with the Co-occurrence method for 2-second OWS, these two codes were the only differences. Since HFI is a frequency-based method, less frequent action units such as *tongue show* (AD19) were not picked up as significant using the HFI method but were still sufficiently distinct to differentiate between stress and no-stress states. The logical interpretation of this pattern is that *tongue show* (AD19) and *lips part* (AU25) are sufficiently distinct to discriminate between stress and neutral states, but absence of the codes cannot exclude stress.

**Fig 3.**
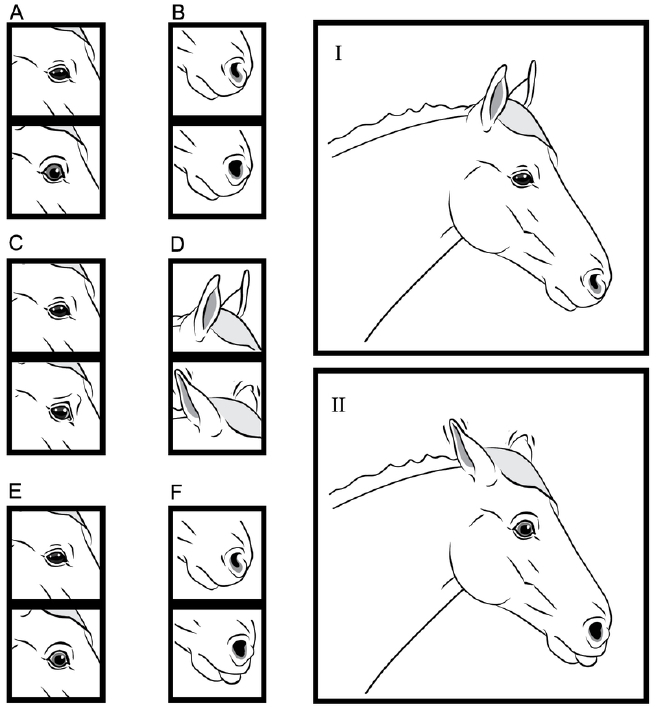
Illustration of facial expressions during stress. Action units (AU)/action descriptors (AD) relevant for recognizing a stressed horse (II). A “neutral” horse (I) is included for reference. A: *Upper lid raiser* (AU5). B: *Nostril dilator* (AD38). C: *Inner brow raiser* (AU101). D: “Ear flicker”. E: *Eye white increase* (AD1). F: *Tongue show* (AD19). Action codes are compared to a “neutral” horse (above). Illustration by Anders Rådén/ARDI.

Most of the codes described by EquiFACS fit well with earlier literature on the subject, e.g., flared nostrils, repetitive mouth behaviors, increased eye white, and an increase in eye movements are features described previously during stress in horses [23,24,30]. However, *inner brow raiser* (AU101) is generally associated with pain and was not expected to be displayed during emotional stress where pain was not present. It is therefore relevant to discuss the presence of other emotional states during the interventions.

Because animals are not able to self-report, biological interpretation of the state of the horses remains open and therefore the meaning of the occurrence of action units remains pragmatic. For example, there is no 100% certainty that the horses are free of pain, since there is no “gold standard” for evaluating pain. However, by only recruiting horses perceived as healthy and free from pain, and by using the horses as their own control, the risk of presence of pain can be lowered, although not completely eliminated. Given this limitation, the overall impressive recall and precision rates of the LOO classification indicate that the action units/action descriptors selected by both the HFI method and the Co-occurrence method can be used successfully to determine whether a horse is stressed or not, on the basis of video-recordings. This is supported by the fact that the methods picked almost identical action units/action descriptors for emotional stress.

### Action descriptors

Since the horse is a flight animal, increased awareness of the surroundings during threats is of great importance to its preservation behaviors. *Dilation of the nostril* (AD38), an effect of several muscles contracting to increase the lumen of the nostril, helps facilitate air intake during flight, meaning that this action descriptor could have a pure physiological purpose, rather than displaying emotion.

The action units *upper lid raiser* (AU5) and *eye white increase* (AD1) increase the field of vision, with the latter being translated mainly into an increase in eye- or head movements. Increased visibility of sclera could also be a result of increased head movements due to restlessness, or of several action units exerting their effects on the eye, e.g., *upper lid raiser* (AU5) or contraction of the infraorbital muscles of the eye, the latter not coded in EquiFACS. This is consistent with earlier findings that a horse under stress tends to focus on the environment and the stressors, causing an increase in head and eye movement [29]. If *eye white increase* (AD1) is present at the same time as *upper lid raiser* (AU5), this could indicate an eye white increase due to raising of the upper eyelid. Otherwise, an increase in *eye white increase* (AD1) could be due to several other factors.

A distinct increase in the movement “ear flicker” was apparent in both transportation and social isolation stress. Ear movements are very communicative [37] and often noted by laymen when describing horse emotions. Movement of the horse’s ears could also aid sound perception, but that effect has been less well studied. During transportation, ear movements due to sound are a likely cause of high “ear flicker” frequency, since the horse’s ear does not linger in any one position for long. During social isolation, a likely cause is increased awareness of the surroundings. The location of the ear conveys information about the horse’s emotional state [6] and is used in many of the grimace scales for pain assessment [10,11]. Only one ear code, *ear rotator* (EAD104), was selected in our dataset, indicating that “ears backward” may be present more often in stressed horses. An increase in “ear flicker” could also prove a good indicator of emotional stress.

### Action units of the upper face

Interesting differences were seen in the action units of the upper face when the two stress induction methods were compared. The reason for *upper lid raiser* (AU5) being more prominently seen during transportation stress, but not selected when analyzing isolation stress, could be that transportation stress is influenced by multiple external factors. In theory, a tension in *m. levator palpebrae superioris* (proposed basis for AU5) would be a plausible mechanism to hide tension in *m. levator anguli occuli medialis* (proposed basis for AU101). This means that it is uncertain whether *inner brow raiser* (AU101) would have been seen more often during transportation if not for the presence of *upper lid raiser* (AU5), due to increased environmental factors or increased intensity of stress. These results suggest that external factors or stress intensity may play a major role in hiding certain facial expressions, meaning that caution is needed in interpretation when the horse is exposed to strong stimuli from the environment (e.g., sounds, smells, visuals).

The frequency of *blink* (AU145) increased during both transportation and social isolation. An earlier study also reported an increase in blinks during stressful situations [16]. However, Merkies et al. [17] found instead that full blink diminished during stress. In this study, the increase was only statistically significant for the Co-occurrence method during transportation stress. This may be a result of the greater number of horses in the transportation stress group. Differences in frequency of full blinks were not significant between baseline and stress (Fig 2). Since duration of full blink is by definition restrained to less than half a second, it is unlikely that maximum duration would have any meaning, even if it showed a significant increase. Blink frequency needs to be studied further in order to draw firm conclusions regarding its role as a marker of stress, and special consideration needs to be given to the induction method.

### Action units of the lower face

The only action unit selected as indicative of stress for the lower face was *lips part* (AU25). Concurrently, increased frequency of *tongue show* (AD19) was noted. This coincides well with earlier findings on behaviors of the tongue and repetitive mouth and licking behaviors during stress [29,35]. These codes were only selected by the Co-occurrence method, but there was an increase in frequency and maximum duration for the codes when tested specifically in the HFI method. As discussed earlier, this might be a result of the codes being less frequent, but highly distinct for stress. *Tongue show* (AD19) and *lips part* (AU25) are similarly interrelated, since *tongue show* (AD19) requires the horse’s mouth to be open. Tongue “twisting” has previously been described as a stereotypic behavior [38]. *Tongue show* (AD19) may be interpreted as a coping mechanism in horses subjected to stress. Oral stereotypies are often reported as a long-term consequence of inability of horses to perform natural behavior, creating chronic stress presenting as oral stereotypies (e.g., cribbing). It is therefore not surprising that *tongue show* (AD19) was less significant during the social isolation stress test, since this intervention is a rather acute form of stress. It is interesting that such coping behaviors were recorded during the transportation stress test, given the relatively short duration of acute stress. Since the increase in heart rate was greater during transportation, the presence of *tongue show* (AD19) might also be a result of higher stress intensity during transportation.

### Signs of stress in pain scales

The specificity of facial expressions across emotional states is of interest for their use as an emotional indicator. To our knowledge, pain is the only emotion to be analyzed to date using EquiFACS and, since pain and stress are intimately connected, comparison of the facial expressions of pain and stress is needed. Rashid *et al.* [27] found that *nostril dilator* (AD38) and *chin raiser* (AU17) were indicative of pain when using both the HFI and Co-occurrence methods. The fact that *nostril dilator* (AD38) is also present during stress may be interpreted in several ways, e.g., it may indicate that pain to some degree also induces emotional stress or that occurrence of this action descriptor is common for either type of intervention. During both stress and pain, respiratory rate of the horse tends to increase, which may be a reason for *nostril dilator* (AD38) being common during both interventions.

Interestingly, some face-based pain scoring tools include facial expressions that were selected here as indicative of emotional pain-free stress. For example, the horse grimace scale [10] includes *ear flattener* (EAD103) and *ear rotator* (EAD104) as elements of the pain scale, while the FAP scale [12] uses eye white increase as an element. The “equine pain face” shows the features “tension of the lower face, rotated ears, dilated nostril and tension above the eye” [11]. All but “tension of the lower face” could be seen in the stressed horses. Furthermore, *upper eyelid raiser* (AU5) could mask presence of “tension above the eye” (here interpreted as AU101) in the “pain face”. Some facial expressions linked to stress can also be found in ethograms based on facial expressions of ridden horses [39], where “exposure of sclera”, “mouth opening and shutting repeatedly”, “ears rotated back or flat”, and “tongue exposed” are present. However, comparison of expressions in ridden and unridden horses require caution, since tack [40] or observers [41] might influence the facial expressions present.

It is highly possible that a horse experiencing pain simultaneously experiences some degree of emotional or physiological stress. When discussing both physiological and behavioral aspects of pain assessment, stress is often described as a complicating factor [13], and use of facial expressions for pain scoring is presented as a “better” approach. Based on our results, the level of stress to which a horse is exposed should be considered during interpretation of facial expressions for pain evaluation. For example in ridden horses, where pain is usually not expected to be present, the predictive values of pain scales may decrease because the level of stress is increased in competition or rider-conflict situations [40], resulting in a stress response being interpreted as pain if the above-mentioned action units are present. This would have consequences for the way in which these horses are treated. A horse experiencing pain is in need of veterinary treatment, while a horse experiencing stress needs the help of behavior specialist or trainer to reduce its emotional stress. EquiFACS coding may be used for further validation of the construct validity of current horse emotion (including pain) assessment tools based on facial expressions [1].

### Using video and EquiFACS to record facial expressions

While obtaining useful footage proved relatively simple, the coding of action units and action descriptors was very time-consuming, with each 30-second sequence taking approximately one hour to annotate manually. In total, all clips took 300 hours of annotation. Based on the good rater agreement, one coding of each sequence appears sufficient.

It is not possible to perform a complete EquiFACS annotation during direct observation, e.g., under clinical conditions. Some easily observable EquiFACS-based measures, such as *increased eye white* (AD1), increased blink rate, and mouth behaviors, should be investigated for their performance value in “grimace”-based scales for discovering stress or no-stress in both undisturbed horses and “disturbed” horses, e.g., ridden horses. As mentioned earlier, many facial features may be affected by external stimuli which may have nothing to do with the emotion of the horse. Therefore, if scorings are not performed directly, we recommend that the analysis be performed on video-recordings, where onset and offset of a certain grimace can be verified, increasing the specificity of that facial action. Still images may capture a short moment where the horse is reacting to other stimuli, such as a noise, that is not detectable from the image. Techniques for automatic detection of action units in humans generally rely on video footage [42]. It has also been shown that, for horses in pain, the chance of picking up all essential action units/descriptors on a random still frame is very low [27]. The same is likely true for emotional stress. If live scorings are used to assess pain, the horse’s emotional state and possible external factors need to be considered when interpreting the results. Many action units are also very difficult to score live, without the use of slow-motion or frame-by-frame in video.

The data presented here provide a foundation for development of automated surveillance of animals based on video-recordings or live video, to determine facial expressions during stress using EquiFACS. However, there were difficulties with the use of both the surveillance cameras and the automated video selection tool. The freely moving stressed horses changed position rapidly, decreasing the length of the optimal observation windows. The automated face detection method used for random and unbiased selection of video clips for annotation tended to prioritize the longest clips, meaning that the clips might have been systematically selected from quiet periods, i.e., from periods where the horse was least stressed. This could be a contributing factor to the lower stress level observed during social isolation than during transportation, and to some action codes not being significant. There were also fewer horses in the social isolation group. The main limitations with this study were the small number of horses and the inability to blind the transportation videos. We sought to overcome this problem by using three annotators for the transportation clips.

## Conclusions

It proved possible to induce and monitor the presence of emotional stress objectively in horses under field conditions, using simple equipment and ordinary management practices. Applying two different frequency and duration-based methods revealed that two types of short-term emotional stress (social isolation, transportation) induced several facial actions. Overall, it was concluded that the facial traits *eye white increase* (AD1), *nostril dilator* (AD38), *inner brow raiser* (AU101), *upper lid raiser* (AU5), “ear flicker*”*, and *tongue show* (AD19) were indicative of equine stress. This confirmed earlier findings of behavioral aspects during stress and some features in equine pain scales. Data from this study can be used to construct less time-consuming observation tools for use when fast scoring is needed (e.g., in an equine hospital environment). Scoring using the EquiFACS system proved to work well using video footage, showing that it can be performed using video surveillance to minimize observer influence and errors during scoring.

## Acknowledgements

DVM Camilla Frisk and DVM Alina Silventoinen are thanked for annotating the films. Horses and owners who participated in this study are warmly thanked for their contributions.

## Supporting information captions

**S1 Dataset. Data used for the data analysis**

**S2 File. HRM files (compressed R-R intervals for heart rate analysis)**

**S3 Video. Sample clip of a baseline video**

**S4 Video. Sample clip of an isolation video**

**S5 Video. Sample clip of a transportation video**

## Reference

1. Descovich KA, Wathan J, Leach MC, Buchanan-Smith HM, Flecknell P, Farningham D, et al. Facial expression: An under-utilized tool for the assessment of welfare in mammals. ALTEX. 2017;34: 409–429. doi:10.14573/altex.1607161

2. Cooke N. A. Facial Mimicry versus Perspective-taking: Decoding Instructional Sets as Empathy-inducing Strategies. M.Sc. Thesis. Appalachian State University. 2015. Available from: https://libres.uncg.edu/ir/asu/listing.aspx?id=18771

3. Diogo R, Wood BA, Aziz MA, Burrows A. On the origin, homologies and evolution of primate facial muscles, with a particular focus on hominoids and a suggested unifying nomenclature for the facial muscles of the Mammalia. J Anat. 2009;215: 300–319. doi:10.1111/j.1469-7580.2009.01111.x

4. Williams AC. Facial expression of pain: An evolutionary account. Behav Brain Sci. 2002;25: 439–455. doi:10.1017/S0140525X02000080

5. Ekman P, Freisen W V, Ancoli S. Facial signs of emotional experience. J Pers Soc Psychol. 1980;39: 1125–1134. doi:10.1037/h0077722

6. Wathan J, Burrows AM, Waller BM, McComb K. EquiFACS: The equine facial action coding system. PLoS One. 2015;10: 1–35. doi:10.1371/journal.pone.0131738

7. Ekman P, Friesen W V., Hager JC. Facial Action Coding System Investigator’s Guide. Psychologist. Salt Lake City, USA: Research Nexus; 2002.

8. Vick SJ, Waller BM, Parr LA, Pasqualini MCS, Bard KA. A cross-species comparison of facial morphology and movement in humans and chimpanzees using the Facial Action Coding System (FACS). J Nonverbal Behav. 2007;31: 1–20. doi:10.1007/s10919-006-0017-z

9. Waller BM, Peirce K, Caeiro CC, Scheider L, Burrows AM, McCune S, et al. Paedomorphic Facial Expressions Give Dogs a Selective Advantage. PLoS One. 2013;8: 1–6. doi:10.1371/journal.pone.0082686

10. Dalla Costa E, Minero M, Lebelt D, Stucke D, Canali E, Leach MC. Development of the Horse Grimace Scale (HGS) as a pain assessment tool in horses undergoing routine castration. PLoS One. 2014;9: 1–10. doi:10.1371/journal.pone.0092281

11. Gleerup KB, Forkman B, Lindegaard C, Andersen PH. An equine pain face. Vet Anaesth Analg. 2015;42: 103–114. doi:10.1111/vaa.12212

12. van Loon JPAM, Van Dierendonck MC. Monitoring acute equine visceral pain with the Equine Utrecht University Scale for Composite Pain Assessment (EQUUS-COMPASS) and the Equine Utrecht University Scale for Facial Assessment of Pain (EQUUS-FAP): A scale-construction study. Vet J. 2015. doi:10.1016/j.tvjl.2015.08.023

13. McLennan KM, Miller AL, Dalla Costa E, Stucke D, Corke MJ, Broom DM, et al. Conceptual and methodological issues relating to pain assessment in mammals: The development and utilisation of pain facial expression scales. Appl Anim Behav Sci. 2019;217: 1–15. doi:10.1016/j.applanim.2019.06.001

14. Mayo LM, Heilig M. In the face of stress: Interpreting individual differences in stress-induced facial expressions. Neurobiol Stress. 2019;10: 100166. doi:10.1016/J.YNSTR.2019.100166

15. Hintze S, Smith S, Patt A, Bachmann I, Würbel H. Are eyes a mirror of the soul? What eye wrinkles reveal about a horse’s emotional state. PLoS One. 2016;11: 1–15. doi:10.1371/journal.pone.0164017

16. Roberts K, Hemmings AJ, Moore-Colyer M, Parker MO, McBride SD. Neural modulators of temperament: A multivariate approach to personality trait identification in the horse. Physiol Behav. 2016;167: 125–131. doi:10.1016/j.physbeh.2016.08.029

17. Merkies K, Ready C, Farkas L, Hodder A. Eye Blink Rates and Eyelid Twitches as a Non-Invasive Measure of Stress in the Domestic Horse. Anim an open access J from MDPI. 2019;9: 562. doi:10.3390/ani9080562

18. Budzyńska M. Stress reactivity and coping in horse adaptation to environment. J Equine Vet Sci. 2014;34: 935–941. doi:10.1016/j.jevs.2014.05.010

19. Schmidt. Cortisol release, heart rate, and heart rate variability in transport-naive horses during repeated road transport. Domest Anim Endocrinol. 2010;39: 205–213. doi:10.1016/J.DOMANIEND.2010.06.002

20. Mal ME, Friend TH, Lay DC, Vogelsang SG, Jenkins OC. Behavioral responses of mares to short-term confinement and social isolation 1. 1991;31: 13–24. doi:10.1016/0168-1591(91)90149-R

21. Buchanan KL. Stress and the evolution of condition-dependent signals. Trends Ecol Evol. 2000;15: 156–160. doi:10.1016/S0169-5347(99)01812-1

22. Butler RK, Finn DP. Stress-induced analgesia. Progress in Neurobiology. 2009. pp. 184–202. doi:10.1016/j.pneurobio.2009.04.003

23. Jennings EM, Okine BN, Roche M, Finn DP. Stress-induced hyperalgesia. Prog Neurobiol. 2014;121: 1–18. doi:10.1016/j.pneurobio.2014.06.003

24. Zolovick A, Upson DW, Eleftheriou BE. Diurnal Variation in Plasma Glukocorticosteroid Levels in the Horse (Equus Caballus). J Endocrinol. 1966;35: 249–253. doi:10.1677/joe.0.0350249

25. Rietmann TR, Stauffacher M, Bernasconi P, Auer JA, Weishaupt MA. The association between heart rate, heart rate variability, endocrine and behavioural pain measures in horses suffering from laminitis. J Vet Med Ser A Physiol Pathol Clin Med. 2004;51: 218–225. doi:10.1111/j.1439-0442.2004.00627.x

26. Hörnicke H, Engelhardt W v., Ehrlein HJ. Effect of exercise on systemic blood pressure and heart rate in horses. Pflügers Arch Eur J Physiol. 1977;372: 95–99. doi:10.1007/BF00582212

27. Rashid M, Silventoinen A, Gleerup KB, Andersen PH. Equine Facial Action Coding System for determination of pain-related facial responses in videos of horses. PLoS One. 2020; Forthcoming

28. Kunz M, Meixner D, Lautenbacher S. Facial muscle movements encoding pain - A systematic review. Pain. 2019;160: 535–549. doi:10.1097/j.pain.0000000000001424

29. Young T, Creighton E, Smith T, Hosie C. A novel scale of behavioural indicators of stress for use with domestic horses. Appl Anim Behav Sci. 2012;140: 33–43. doi:10.1016/j.applanim.2012.05.008

30. Tateo A, Padalino B, Boccaccio M, Maggiolino A, Centoducati P. Transport stress in horses: Effects of two different distances. J Vet Behav Clin Appl Res. 2012;7: 33–42. doi:10.1016/j.jveb.2011.04.007

31. Rietmann TR, Stuart AEA, Bernasconi P, Stauffacher M, Auer JA, Weishaupt MA. Assessment of mental stress in warmblood horses: Heart rate variability in comparison to heart rate and selected behavioural parameters. Appl Anim Behav Sci. 2004. doi:10.1016/j.applanim.2004.02.016

32. Rashid M, Broome S, Andersen PH, Gleerup KB, Lee YJ. What should I annotate? An automatic tool for finding video segments for EquiFACS annotation. In: A.J. Spink, editor. Measuring Beaviour. Manchester; 2018. pp. 6–8.

33. ELAN (Version 5.4). Nijmegen: Max Planck Institute for Psycholinguistics; Available: https://tla.mpi.nl/tools/tla-tools/elan/

34. Varoquaux G, Buitinck L, Louppe G, Grisel O, Pedregosa F, Mueller A. Scikit-learn: Macine Learning in Python. GetMobile Mob Comput Commun. 2015;19: 29–33. doi:10.1145/2786984.2786995

35. Munsters CCBM, de Gooijer J-W, van den Broek J, van Oldruitenborgh-Oosterbaan MMS. Heart rate, heart rate variability and behaviour of horses during air transport. Vet Rec. 2013;172: 15–15. doi:10.1136/vr.100952

36. Yngvesson J, de Boussard E, Larsson M, Lundberg A. Loading horses (Equus caballus) onto trailers—Behaviour of horses and horse owners during loading and habituating. Appl Anim Behav Sci. 2016;184: 59–65. doi:10.1016/j.applanim.2016.08.008

37. Wathan J, McComb K. The eyes and ears are visual indicators of attention in domestic horses. Curr Biol. 2014;24: R677–R679. doi:10.1016/j.cub.2014.06.023

38. Fureix C, Gorecka-Bruzda A, Gautier E, Hausberger M. Cooccurrence of Yawning and Stereotypic Behaviour in Horses (Equus caballus). ISRN Zool. 2011: 1–10. doi:10.5402/2011/271209

39. Dyson S, Berger J, Ellis AD, Mullard J. Development of an ethogram for a pain scoring system in ridden horses and its application to determine the presence of musculoskeletal pain. J Vet Behav Clin Appl Res. 2018. doi:10.1016/j.jveb.2017.10.008

40. Gleerup KB, Andersen PH, Wathan J. What information might be in the facial expressions of ridden horses? Adaptation of behavioral research methodologies in a new field. Journal of Veterinary Behavior: Clinical Applications and Research. 2018. doi:10.1016/j.jveb.2017.12.002

41. Torcivia C, Mcdonnell S. In-Person Caretaker Visits Disrupt Ongoing Discomfort Behavior in Hospitalized Equine Orthopedic Surgical Patients. Animals. 2020;10. doi:10.3390/ani10020210

42. Littlewort G, Bartlett MS, Fasel I, Susskind J, Movellan J. Dynamics of facial expression extracted automatically from video. Image Vis Comput. 2006;24: 615–625. doi:10.1016/j.imavis.2005.09.011

